# *ddcP, pstB*, and excess D-lactate impact synergism between vancomycin and chlorhexidine against *Enterococcus faecium* 1,231,410

**DOI:** 10.1101/450593

**Authors:** Pooja Bhardwaj, Moutusee Z. Islam, Christi Kim, Uyen Thy Nguyen, Kelli L. Palmer

## Abstract

Vancomycin-resistant enterococci (VRE) are important nosocomial pathogens that cause life-threatening infections. To control hospital-associated infections, skin antisepsis and bathing utilizing chlorhexidine is recommended for VRE patients in acute care hospitals. Previously, we reported that exposure to inhibitory chlorhexidine levels induced the expression of vancomycin resistance genes in VanA-type *Enterococcus faecium*. However, vancomycin susceptibility actually increased for VanA-type *E. faecium* in the presence of chlorhexidine. Hence, a synergistic effect of the two antimicrobials was observed. In this study, we used multiple approaches to investigate the mechanism of synergism between chlorhexidine and vancomycin in the VanA-type VRE strain *E. faecium* 1,231,410. We generated clean deletions of 7 of 11 *pbp*, transpeptidase, and carboxypeptidase genes in this strain (*ponA, pbpF, pbpZ, pbpA, ddcP, ldt*_fm_, and *vanY*). Deletion of *ddcP*, encoding a membrane-bound carboxypeptidase, altered the synergism phenotype. Furthermore, using *in vitro* evolution, we isolated a spontaneous synergy escaper mutant and utilized whole genome sequencing to determine that a mutation in *pstB*, encoding an ATPase of phosphate-specific transporters, also altered synergism. Finally, addition of excess D-lactate, but not D-alanine, enhanced synergism. Overall, our work identified factors that alter chlorhexidine-induced vancomycin resensitization in a model VanA-type VRE strain.

## Introduction

*Enterococcus faecium* are Gram-positive commensal bacteria inhabiting the gastrointestinal tracts of humans and animals (1). The ability to survive in harsh environmental conditions including starvation and desiccation facilitated the emergence of hospital-adapted strains which are resistant to the action of antibiotics and disinfectants (2). Hospital-adapted enterococcal strains have limited treatment options and are typically characterized by high-level resistance to vancomycin, a glycopeptide antibiotic which inhibits the process of peptidoglycan synthesis (3, 4). Vancomycin-resistant enterococci (VRE) synthesize peptidoglycan precursors for which vancomycin has low affinity (5-8). Vancomycin resistance in hospital-adapted enterococcal isolates occurs through the horizontal acquisition of resistance genes (9, 10). For VanA-type VRE, vancomycin resistance is conferred and controlled by the activities encoded by the *vanRS, vanHAX*, and *vanYZ* genes.

Patients in critical care units are frequently bathed or cleansed with chlorhexidine, a cationic cell membrane-targeting antimicrobial, to reduce the occurrence of hospital-associated infections (11-13). Chlorhexidine interacts with the negatively charged phospholipids and proteins on the cell membrane after primary adsorption by the cell (14, 15). Low chlorhexidine levels disrupt the membrane potential and integrity whereas high chlorhexidine levels can cause a complete precipitation of the cytoplasm (16-18). We previously analyzed the transcriptome of the VanA-type vancomycin-resistant strain *E. faecium* 1,231,410 exposed to inhibitory levels of chlorhexidine, and we found that chlorhexidine stress induced the expression of the VanA-type vancomycin resistance genes (19). However, vancomycin MIC actually decreased when chlorhexidine was present in broth microdilution assays (19).

We previously proposed three models to explain chlorhexidine-induced vancomycin sensitization despite transcriptional activation of VanA-type vancomycin resistance genes by chlorhexidine (19). Vancomycin resistance genes code for the synthesis of alternative peptidoglycan precursors that terminate in D-alanine-D-lactate (D-Ala-D-Lac), for which vancomycin has lower affinity compared to the normal D-alanine-D-alanine (D-Ala-D-Ala). Model 1 is that altered penicillin-binding protein (Pbp) levels in the presence of chlorhexidine prevents D-Ala-D-Lac precursors from being cross-linked. Model 2 proposes that chlorhexidine alters substrate pools for peptidoglycan synthesis, resulting in vancomycin-sensitive termini that are neither D-Ala-D-Ala nor D-Ala-D-Lac. Model 3 is that post-translational regulation of VanX and/or VanY prevents depletion of D-Ala-D-Ala termini from peptidoglycan precursors in the presence of chlorhexidine. In this study, we used targeted gene deletions, *in vitro* evolution, and culture assays in modified media to assess specific features of these models. Overall, we identify two genes, *ddcP* and *pstB*, and one growth condition (excess D-lactate), that alter chlorhexidine-induced vancomycin sensitization in the model VanA-type VRE strain *E. faecium* 1,231,410.

## Materials and Methods

### Bacterial strains and growth conditions

Bacterial strains used in this study are shown in Table 1. *E. faecium* was cultured at 37°C on brain heart infusion (BHI) agar or in broth without agitation unless otherwise stated. *Escherichia coli* was cultured at 37°C in lysogeny broth (LB) broth with shaking at 225 rpm or on LB with 1.5% agar unless otherwise stated. The chlorhexidine product used for all experiments was Hibiclens (4% wt/vol chlorhexidine gluconate with 4% isopropyl alcohol). We refer to Hibiclens as H-CHG in this study. Antibiotics were added at the following concentrations: vancomycin, 50 μg/ml for *E. faecium*, and chloramphenicol, 15 μg/ml for *E. coli*.

**Table 1.**
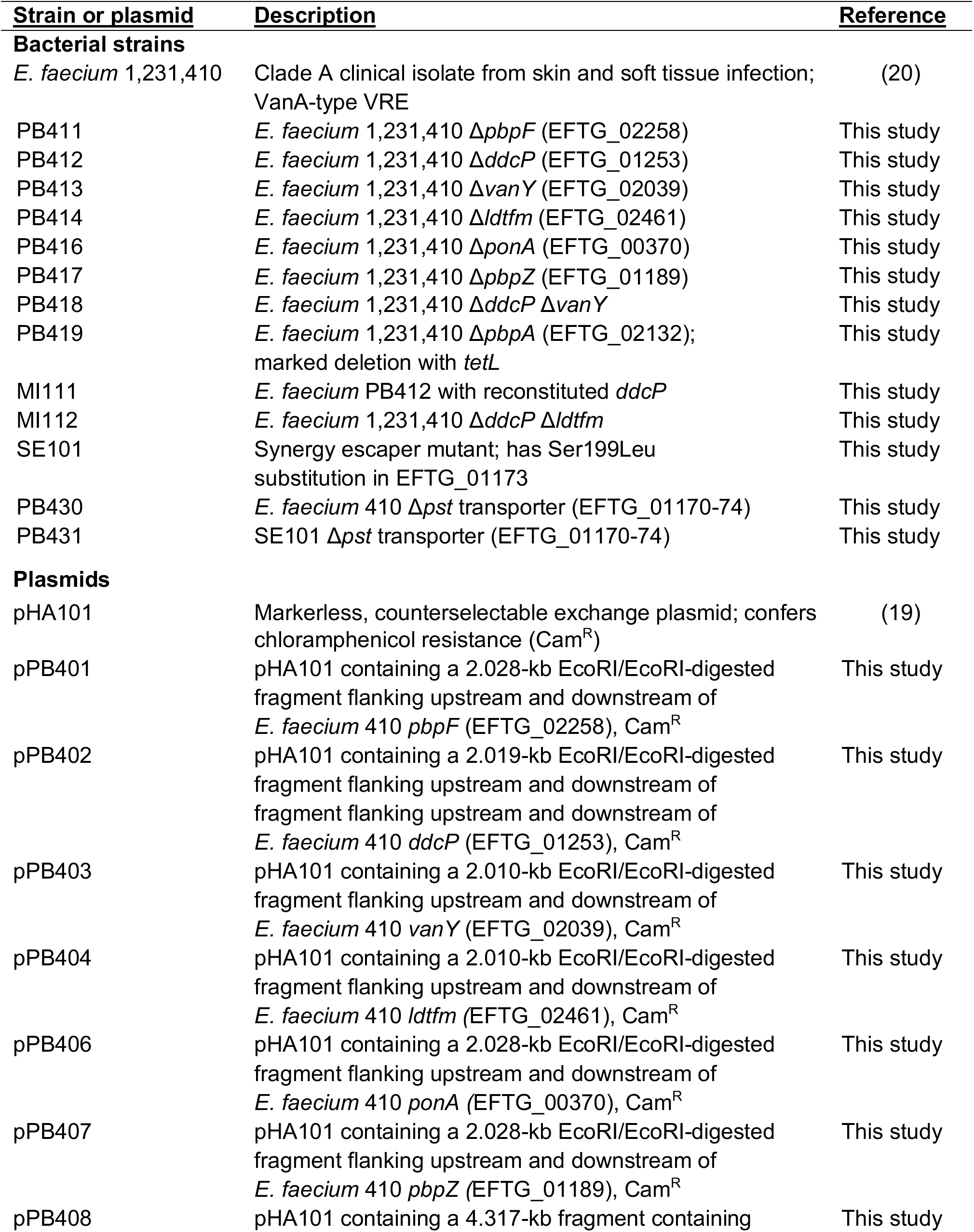
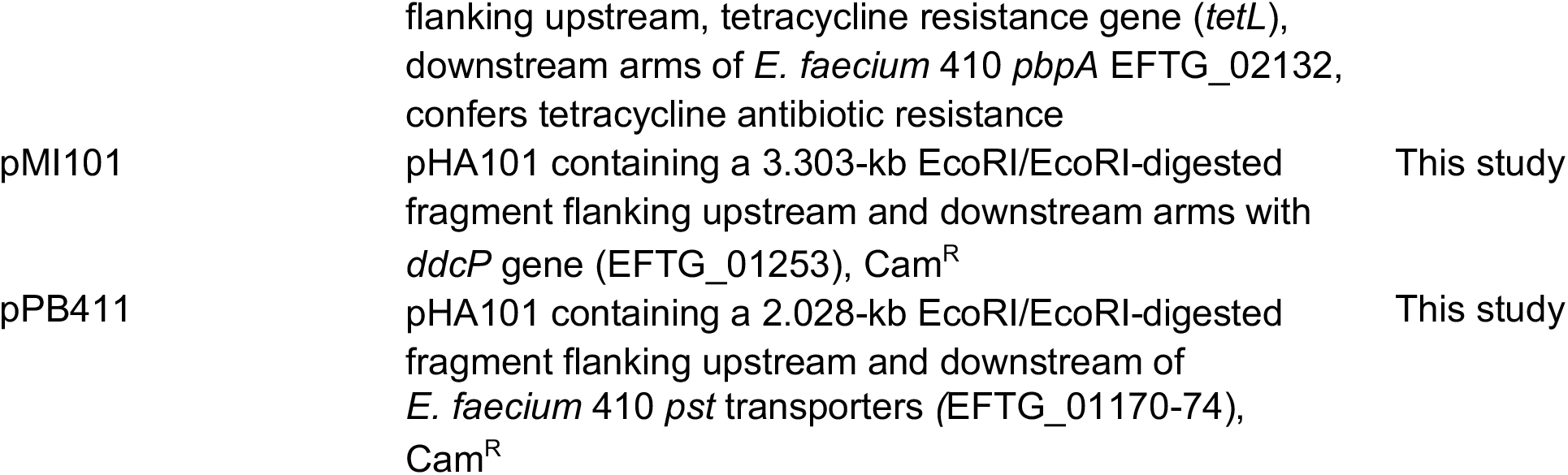
Bacterial strains and plasmids used in the study.

### Routine molecular biol ogy techniques

*E. faecium* genomic DNA (gDNA) was isolated using a previously published protocol (21). Electroporation of *E. faecium* was performed as described previously (19). Plasmid s were purified using the GeneJET Miniprep kit (Thermo Scientific). DNA fragments were purified using the Purelink PCR purification kit (Invitrogen). *Taq* polymerase (New England Biolabs; NEB) was used for routine PCR reactions. Phusion polymerase (Fisher) was used for cloning applications. Restriction endonucleases (NEB) and T4 DNA ligase (NEB) reactions were performed per the manufacturer’s instructions. Routine DNA sequencing was performed by the Massachusetts General Hospital DNA core facility (Boston, MA). All genetic constructs were validated by DNA sequencing. Primers used in this study are shown in Table S1.

### MIC determinations

The minimum inhibitory concentration (MIC) was defined as the lowest drug concentration at which the OD_600nm_ of the bacterial culture matched the OD_600nm_ of the negative control (uninoculated BHI broth). For this study, we refer to synergy MIC as the vancomycin MIC of enterococci in BHI supplemented with H-CHG. The synergy MIC was determined by slightly modifying our previously published protocol (19). 5 μl of vancomycin stock solution (40 mg/ml in water) was added to 195 μl of BHI supplemented with H-CHG in the first well of a 96-well microtiter plate. Next, 100 μl was transferred to the next well containing 100 μl of BHI supplemented with H-CHG to make two-fold serial dilutions of vancomycin drug. Overnight cultures of *E. faecium* were diluted to OD_600_ of 0.01 in fresh BHI, and 5 μl of the diluted culture was used to inoculate the wells of the plate. The OD_600_ of the cultures was measured after 24 h incubation at 37°C. For determining synergy MIC in the presence of D-lactate or D-alanine, D-lactate or D-alanine were solubilized to a final concentration of 0.2 M in BHI and the solutions were filter sterilized. Two-fold serial dilutions of vancomycin were made in BHI supplemented with D-lactate/D-alanine and H-CHG as described above. Fold decrease was calculated by dividing the vancomycin MIC in the absence of H-CHG by the vancomycin MIC in the presence of the highest H-CHG concentration at which visible growth was observed. Each experiment was performed independently at least three times.

### Deletion of genes in *E. faecium*

Genes were deleted in-frame utilizing plasmid pHA101 as described previously (19). Briefly, <1 kb regions upstream and downstream of the target gene were amplified and ligated to pHA101. The sequence of the deletion construct plasmid was verified by Sanger sequencing and introduced into *E. faecium* by electroporation. Temperature shifting at the non-permissible temperature of 42°C and counter-selection with *p*-chlorophenylalanine was followed according to a previously published protocol (22). Presumptive deletion mutants were confirmed by Sanger sequencing of the region of interest followed by Illumina genome sequencing (see below).

After several unsuccessful attempts to generate an unmarked deletion of *pbpA* in *E. faecium* 1,231,410, we introduced a tetracycline resistance marker (*tetL*) between the flanking upstream and downstream arms of the deletion construct to select for deletion mutants. Briefly, *tetL* was amplified from pLT06-*tet* using primers *tetL* For and Rev. The deletion construct was linearized via PCR using Phusion DNA polymerase (Fisher) and primers *pbpA*-linear For and Rev (Table S1). The linearized PCR product was dephosphorylated using Shrimp Alkaline phosphatase (New England Biolabs) per the manufacturer’s instructions and then ligated with *tetL* to generate the deletion construct PB408. The deletion construct was propagated in EC1000 and sequenced using Sanger sequencing prior to transformation into *E. faecium* 1,231,410.

### Complementation of *ddcP* deletion

The *ddcP* gene was restored to the chromosome of the *E. faecium* 1,231,410 *ddcP* deletion mutant. The *ddcP* gene and <1 kb regions up- and downstream were amplified from *E. faecium* 1,231,410 wild-type gDNA, and the amplicon was digested and inserted into pHA101. The knock-in plasmid construct (pMI101) was transformed into the *ddcP* deletion mutant by electroporation. The temperature shifting and counter-selection protocol was followed as described previously (22). The chromosomal integration of the gene was confirmed by Sanger sequencing.

### Growth kinetics of *E. faecium* in the presence of H-CHG and vancomycin

Overnight cultures of *E. faecium* were diluted to an OD_600_ of 0.01 in fresh, pre-warmed BHI and incubated at 37°C with shaking at 100 rpm. The cultures were grown until OD_600_ reached 0.5 to 0.6. Twenty-five milliliters of the culture were added to equal amounts of pre-warmed BHI containing vancomycin (50 μg/ml) and/or H-CHG (4.9 μg/ml), or only BHI (control). OD_600_ values were then monitored hourly for 6 h, and an OD_600_ reading was taken at the 24 h time point. For some experiments and timepoints, CFU/mL were additionally determined by serial dilution of culture and plating on BHI agar. Growth curves were repeated independently three times. For assessing synergy between vancomycin and glycine, the same experimental design was used, except that H-CHG was replaced with 0.2 M glycine.

### Isolation of *E. faecium* 1,231,410 synergy escaper mutant

An *E. faecium* 1,231,410 wild-type culture treated with vancomycin and H-CHG was incubated for 24 h, when turbidity was observed. The recovered culture was used as an inoculum for a second growth curve experiment with vancomycin and H-CHG. OD_600_ values were monitored for 6 h, and at the end of 6 h, the cultures were cryopreserved at −80°C. The stocked populations were struck on BHI agar, and the synergy MIC was determined for well-isolated colonies using the broth microdilution assay described above. Colonies with elevated synergy MIC as compared to the parental *E. faecium* 1,231,410 strain were passaged three times in BHI broth and the synergy MICs determined again. A strain with a stably elevated synergy MIC was isolated; this strain is referred to as SE101.

### Genome sequencing and analysis

SE101 gDNA was isolated according to a previously published protocol (21) and sequenced with Illumina technology at Molecular Research LP (Shallowater, Texas). Paired end, 2×150 reads were obtained. For the analyses, sequence reads were assembled locally to the *E. faecium* 1,231,410 draft reference genome (GenBank accession number NZ_ACBA00000000.1) using default parameters in CLC Genomics Workbench (Qiagen). Polymorphisms in the resequencing assemblies were detected using basic variant mapping using default settings with a minimum variant frequency of 50%. To detect transposon/IS element hopping, the assembly parameters were changed to global instead of local alignment, and regions with sequential polymorphisms were manually analyzed for potential transposon/IS element hopping. Sanger sequencing was utilized to confirm polymorphisms.

To confirm deletion mutants, gDNA was isolated as above and sequenced with Illumina technology at the UT-Dallas Genome Core (for deletion mutants Δ*pbpF*, Δ*ddcP*, Δ*vanY*, Δ*ldtfm*, Δ*ponA*, Δ*ddcP* Δ*vanY*, Δ*ddcP* Δ*ldtfm*, and the Δ*pbpA* marked deletion with *tetL*) or the Microbial Genome Sequencing Center in Pittsburgh, PA (for the Δ*pbpZ* mutant). Paired end, 2×75 reads and 2×150 reads were obtained, respectively. For the analyses, sequence reads were assembled locally to the *E. faecium* 1,231,410 draft reference genome as above. Average coverage of the reference genome ranged from 88-245X across all mutants analyzed. The absence of the gene of interest was confirmed for each presumptive deletion mutant by manually analyzing the read assembly at the location of the gene.

### Phosphate levels measurement

A commercially available kit (Sigma MAK030) and previously published protocol (23, 24) was utilized to measure intracellular inorganic phosphate (Pi) levels at five time points (OD_600_ from 0.4-0.5, 0.6-0.7, 0.7-0.8, 0.8-0.9, 1.0-1.5) from *E. faecium* 1,231,410 and SE101 cultures. The phosphate levels were normalized by CFU and five independent trials were performed.

### Accession number

Raw Illumina sequencing reads generated in this study are available in the Sequence Read Archive under the accession numbers SRP113791 (for strain SE101) and PRJNA657813 (for confirmation of deletion mutants).

## Results and Discussion

### Addition of D-lactate enhances chlorhexidine-induced vancomycin sensitization

It was previously observed that culture supplementation with 0.2 M of glycine or select D-amino acids (including D-methionine, D-serine, D-alanine, or D-phenylalanine) increased VRE susceptibility to vancomycin (25). Consistent with this, Aart et al reported that excess D-Ala substrate competes with D-Lac, thereby increasing the ratio of cell wall termini ending at D-Ala and the efficacy of vancomycin against *Streptomyces* and *E. faecium* (26). We reasoned that if H-CHG stress resulted in an alteration of substrate pools and therefore vancomycin-sensitive termini that are neither D-Ala-D-Ala nor D-Ala-D-Lac (Model 2), an excess of D-lactate could compete with this alternative pathway, thereby increasing the number of D-Ala-D-Lac termini and resulting in loss of synergism between vancomycin and H-CHG.

To test this, synergy assays were performed with *E. faecium* 1,231,410 in the presence of 0.2 M D-lactate or D-alanine (Table 2). Addition of D-lactate to BHI broth lacking H-CHG resulted in a 4-fold increase in vancomycin MIC (Table 2), demonstrating that excess D-lactate does result in reduced vancomycin susceptibility. However, and counter to our expectations, in the presence of both D-lactate and H-CHG, the H-CHG-induced vancomycin resensitization phenotype was enhanced (Table 2). This result suggests that the synergism phenotype is dependent upon D-Ala-D-Lac termini and is enhanced by increased abundance of D-Ala-D-Lac termini. As expected based on the results of Aart et al (26), vancomycin MIC decreased in the presence of D-alanine, and we observed only a 2-fold additional MIC decrease in the presence of both D-alanine and H-CHG (Table 2).

**Table 2.** Median vancomycin MICs in *E. faecium* 1,231,410.

|  | Vancomycin MIC (µg/ml)<sup>a</sup> |  |  |
| --- | --- | --- | --- |
| H-CHG (µg/ml)<sup>b</sup> | BHI | BHI + 0.2 M D-Lactate | BHI + 0.2 M D-Alanine |
| 0 | 250 | 1000 | 7.8 |
| 1.2 | 62.5 | 1.0 | 3.9 |
| 2.4 | 0.2 | No growth | No growth |
| 4.9 | No growth | No growth | No growth |

<sup>a</sup>Vancomycin MICs (µg/ml) at 24 h post-inoculation from at least three independent experiments.
<sup>b</sup>4.9 µg/ml H-CHG is the MIC for *E. faecium* 1,231,410; 1.2 µg/ml H-CHG and 2.4 µg/ml H-CHG are 1/4X MIC and 1/2X MIC, respectively.

### Deletion of *ddcP* alters the synergism phenotype

The enterococcal cell wall is a multi-layered network and is characterized by the presence of peptidoglycan, teichoic acid, and polysaccharides (27, 28). The main component of the cell wall is peptidoglycan, which is a mesh-like structure consisting of parallel glycan chains cross-linked by amino acids (27). The glycan chains consist of two alternating amino sugars, N-acetylglucosamine (GlcNAc) and N-acetylmuramic acid (MurNAc), connected by β-1,4 linkages (29-31). In *E. faecium*, each MurNAc sugar is linked to short stem pentapeptides (L-alanine^1^-D-isoglutamic acid^2^-L-lysine^3^-D-alanine^4^-D-alanine^5^), which alternate between L- and D-amino acids (30, 32). The MurNAc and GlcNAc glycan sugars are synthesized as a UDP (Uridine diphosphate) derivative in a step-wise fashion in the cytosol (33). Next, MurNAc sugars containing short peptides are transferred to a lipid carrier Lipid I (C55-undecaprenol) (34, 35) and added to UDP-derivative GlcNAc to build a disaccharide, GlcNAc-MurNAc-pentapeptide-C55 pyrophosphate, also known as Lipid II (33, 34). Lipid II units are translocated from the cytosol to the outer side of the cell membrane (36) and polymerized through an ordered rate of two processes, transglycosylation (condensation of linear glycan chains) and transpeptidation (cross-linking between carboxyl group of one pentapeptide and amino acid of an adjacent pentapeptide). The disaccharide units are integrated into the growing peptidoglycan layers to form the cell wall (37, 38) and the lipid carrier is recycled back into the cytosol.

Two classes of penicillin-binding proteins (Pbps) mediate the transpeptidation process (39-41). Class A Pbps (encoded by *ponA, pbpF*, and *pbpZ*) are bifunctional, multimodular, high-molecular mass proteins, and catalyze both transpeptidation and transglycosylation reactions.

Class B Pbps (encoded by *pbpB, pbpA*, and *pbp5*) are monofunctional, low-molecular mass proteins, and catalyze only transpeptidation reactions. Class A and B Pbps mediate 4,3 cross-links (D,D-transpeptidation) between cell wall precursors and these cross-links constitute the majority of the mature cell wall (38). However, 3,3 cross-links are also present in the enterococcal cell wall. The combined activities of DdcP or DdcY (D,D-carboxypeptidase) and the L,D transpeptidase Ldt_fm_ can bypass conventional D,D-transpeptidation and mediate 3,3 cross-linking (42-44). DdcP and/or DdcY generates tetrapeptides and reduces the availability of pentapeptide precursors by trimming the terminal D-Ala. Next, Ldt_fm_ mediates cross-links between these cell wall termini.

To determine whether these factors contribute to vancomycin-chlorhexidine synergism against VRE (Model 1), we deleted 6 of 9 of these genes in *E. faecium* 1,231,410 (Table 1). Our presumptive *pbpB, pbp5*, and *ddcY* deletion mutants could not be confirmed by genome sequencing. We utilized growth curves to assess phenotypes of the deletion mutants in the presence and absence of vancomycin and H-CHG. For these experiments, vancomycin (50 μg/ml) and H-CHG (4.9 μg/ml) were added to exponentially growing *E. faecium* cultures; a no-drug control was also performed. As shown in Fig. 1A, the OD_600_ of *E. faecium* 1,231,410 wild-type cultures decreased after addition of vancomycin and H-CHG, consistent with cell lysis. After 24 h, the cultures treated with vancomycin and H-CHG were visibly turbid, indicating that *E. faecium* can recover from the effects of the antimicrobials in this experimental condition.

**Figure 1.**
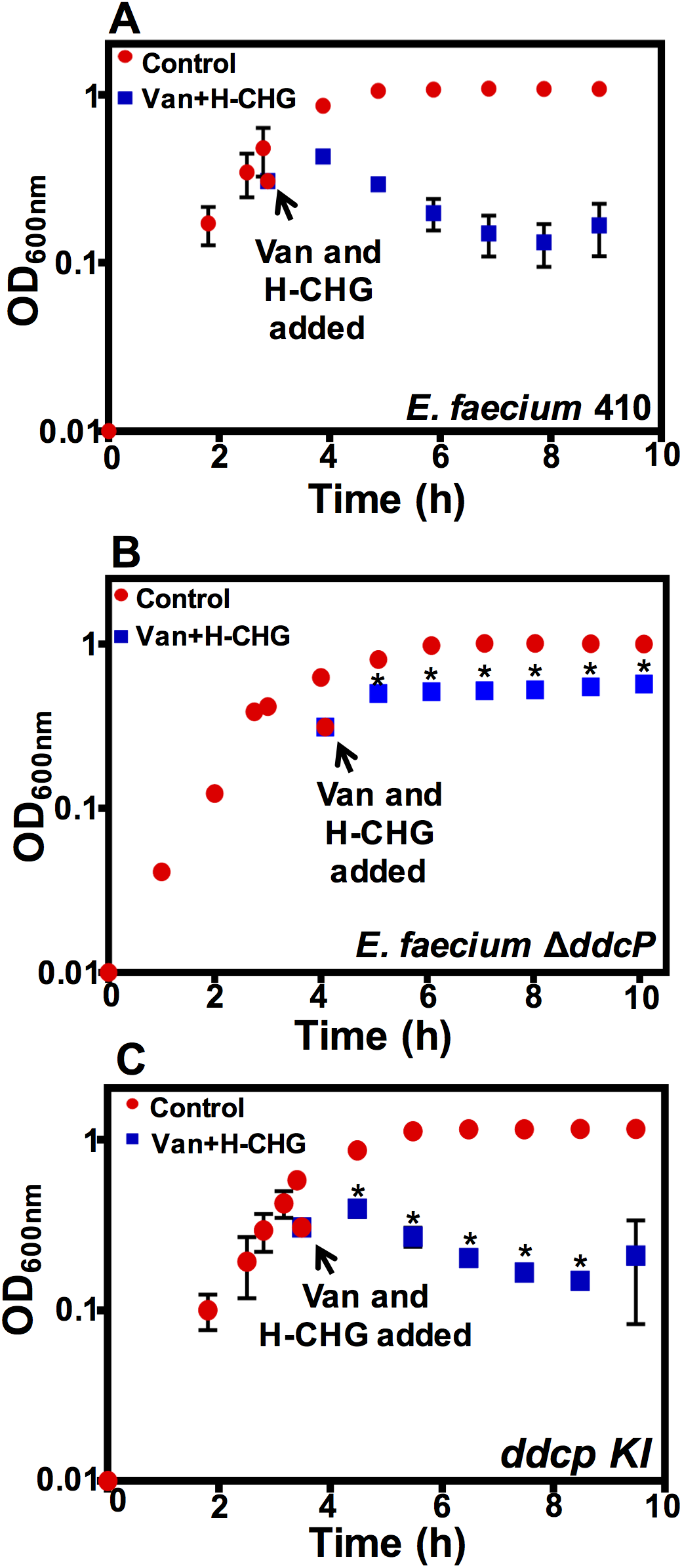
Δ*ddcP* mutant has an altered vancomycin/H-CHG synergy phenotype. Optical density (OD_600nm_) of (A) *E. faecium* 1,231,410 wild-type (*E. faecium* 410), (B) the *ddcP* deletion mutant, and (C) the *ddcP* complemented strain with and without vancomycin and H-CHG treatment. *E. faecium* was cultured in BHI broth until the OD_600_ reached 0.6, as described in materials and methods. Equal volumes of cultures were split into BHI (control; red circles) or BHI containing vancomycin (50 μg/ml) and H-CHG (4.9 μg/ml) (blue squares). OD_600_ values were monitored for 6 h. Error bars indicate standard deviations from n=3 independent experiments. Significance was assessed using the one-tailed Student’s *t*-test. * denotes *P*-value < 0.05. Stars indicate significant differences between vancomycin- and H-CHG-treated cultures in panel B versus A, and in panel C versus B. Note that growth curve of *E. faecium* 410 wild-type in the presence of vancomycin and chlorhexidine from Fig. 1A has been shown again in Fig. S1A for comparison with the *pbp* deletion mutants.

The growth phenotypes of the Δ*pbpF*, Δ*ponA*, Δ*pbpZ*, Δ*pbpA*, and Δ*ldt_fm_* deletion mutants, as measured by OD_600_ values, were comparable to the parental strain (Fig. S1). The Δ*ddcP* mutant had a different phenotype from the wild-type strain (Fig. 1B). After treatment with vancomycin and H-CHG, the OD_600_ values for the *ddcP* deletion mutant did not decrease. The OD_600_ values were significantly different between the wild-type and the Δ*ddcP* mutant for all time points post-H-CHG and vancomycin addition (*P*-value < 0.05, one-tailed Student’s *t* test). The Δ*ddcP* mutant was complemented by restoration of the *ddcP* gene in *cis*. OD_600_ values of the complemented strain in the presence of vancomycin and H-CHG were similar to the wild-type (Fig. 1C). No statistically significant differences in OD_600_ values were observed between the wild-type and the *ddcP* complemented strain in the presence of vancomycin and H-CHG.

In separate experiments, we assessed both the OD_600_ and viability of wild-type and D*ddcP* cultures for 3 hours post-treatment with vancomycin and H-CHG. Optical density values significantly differed between the two strains post-treatment; however, CFU/mL values did not (Fig. S2). These results suggest that *ddcP* deletion does not alter the mechanism by which vancomycin/H-CHG kills *E. faecium*, but rather whether the dead cells lyse.

### The Δ*ddcP* mutant has altered susceptibility to vancomycin-glycine synergy

Synergism between glycine and vancomycin was previously reported for VRE (25). We carried out growth curves in the presence of vancomycin (50 μg/ml) and 0.2 M glycine. A lytic effect was observed for the wild-type strain cultured with vancomycin and glycine, with the most pronounced effect observed at the 24 h time point (Fig. 2A). However, this lytic effect was not observed for the Δ*ddcP* mutant (*P*-value < 0.05, assessed by one-tailed Student’s *t*-test) (Fig. 2B). Together with our results in Figure 1 and Figure S2, these results indicate that DdcP activity weakens the cell wall under antimicrobial stress by reducing the availability of cell wall pentapeptide precursors, contributing to cell lysis. This suggests that *ddcP* expression in cells in the presence of CHX and vancomycin antimicrobials contributes to increased cell lysis and synergism between the two antimicrobials.

**Figure 2.**
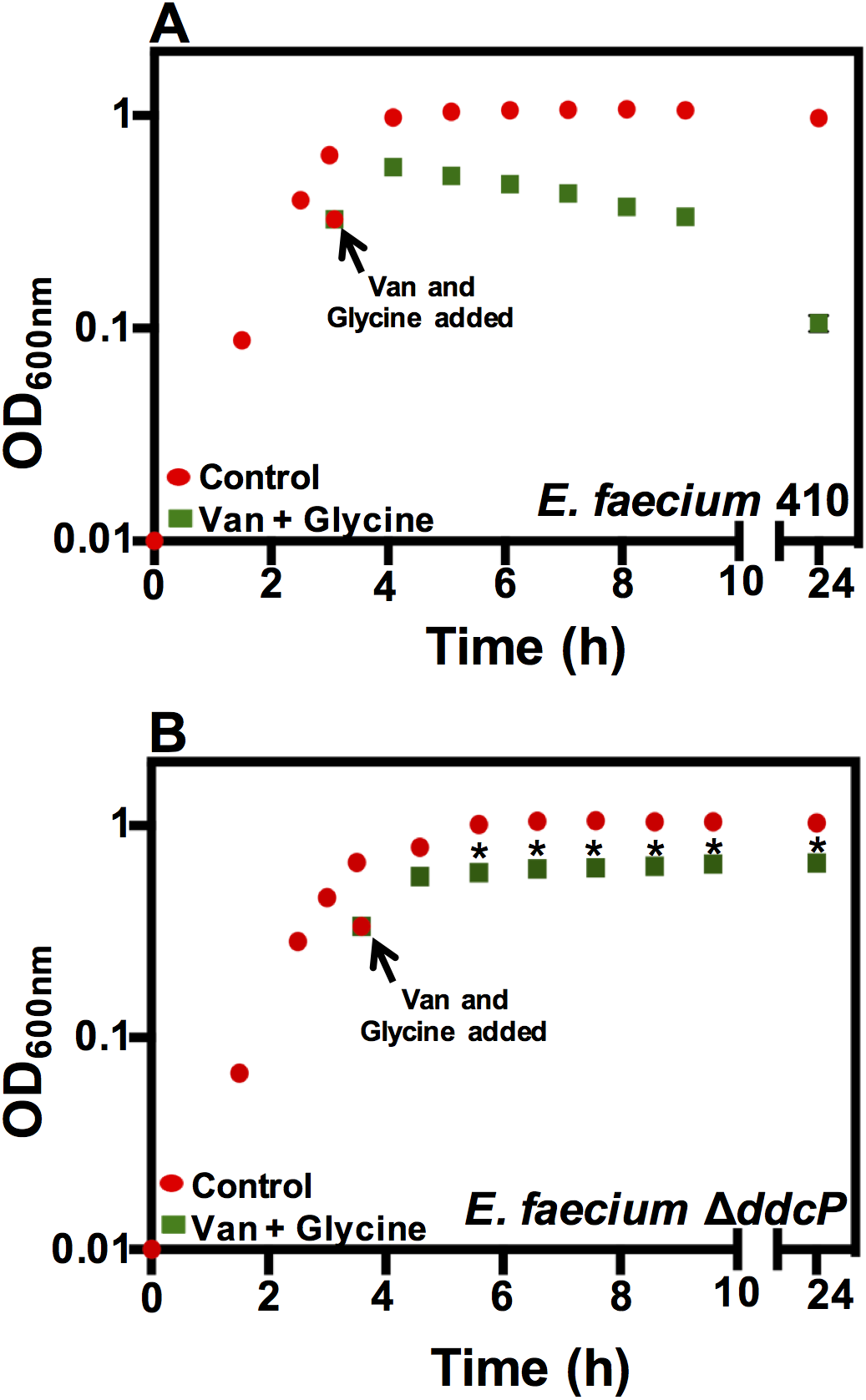
A Δ*ddcP* mutant has reduced susceptibility to vancomycin/glycine synergy. (A) *E. faecium* 410 wild-type and (B) *ddcP* deletion mutant cultures were grown at 37°C in BHI until OD_600_ reached 0.6 as described in materials and methods. Equal volumes of cultures were split into BHI (control; red circles) or BHI containing vancomycin (50 μg/ml) and glycine (0.2 M) (green squares). OD_600_ values were monitored for 6 h and a reading at 24 h was recorded. Error bars indicate standard deviations from n=3 independent experiments. Significance was assessed using the one-tailed Student’s *t*-test. * denotes *P*-value < 0.05. Stars indicate significant differences between vancomycin- and glycine-treated cultures in panel B versus A.

### A *vanY_A_* mutant does not have an altered synergy phenotype

Previously, Kristich et al investigated the genetic basis of synergism between vancomycin and cephalosporins (a class of β-lactam antibiotics) in the VanB-type VRE strain *E. faecalis* V583 (45). The synergism was mediated by VanY_B_, a carboxypeptidase that reduces the availability of precursors ending at D-Ala-D-Ala by trimming the terminal D-Ala, thereby eliminating the target of vancomycin. In the absence of *vanY_B_*, cross-linking of cell wall precursors was mediated by low-affinity Pbps and synergism between vancomycin and cephalosporins was lost (45). To determine whether *vanY_A_* contributed to vancomycin-chlorhexidine synergism against VRE, which is a component of our Model 3, we deleted *vanY_A_* in *E. faecium* 1,231,410. We observed no effect on the synergy phenotype as assessed by OD_600_ values (Fig. S1C). Moreover, deletion of *ddcP* in a Δ*vanY* background did not further enhance the phenotype of a Δ*ddcP* mutant (Fig. S2E).

### *E. faecium* 1,231,410 can escape from vancomycin-chlorhexidine synergy

We observed the growth kinetics of *E. faecium* 1,231,410 cultures exposed to no, either, or both 50 μg/ml vancomycin and 4.9 μg/ml H-CHG over a two-day growth curve experiment. As shown in Fig. 3A, cultures exposed to vancomycin were growth-inhibited for the first 2.5 h after exposure, and after 2.5 h, OD_600_ began to increase, consistent with the induction of vancomycin resistance genes and synthesis of modified cell walls, as previously observed (46, 47). The cultures exposed to H-CHG were also temporarily growth-inhibited. Consistent with the experiments shown in Fig. 1A, the OD_600_ of cultures exposed to both vancomycin and H-CHG declined, and after 24 h, the cultures recovered.

**Figure 3.**
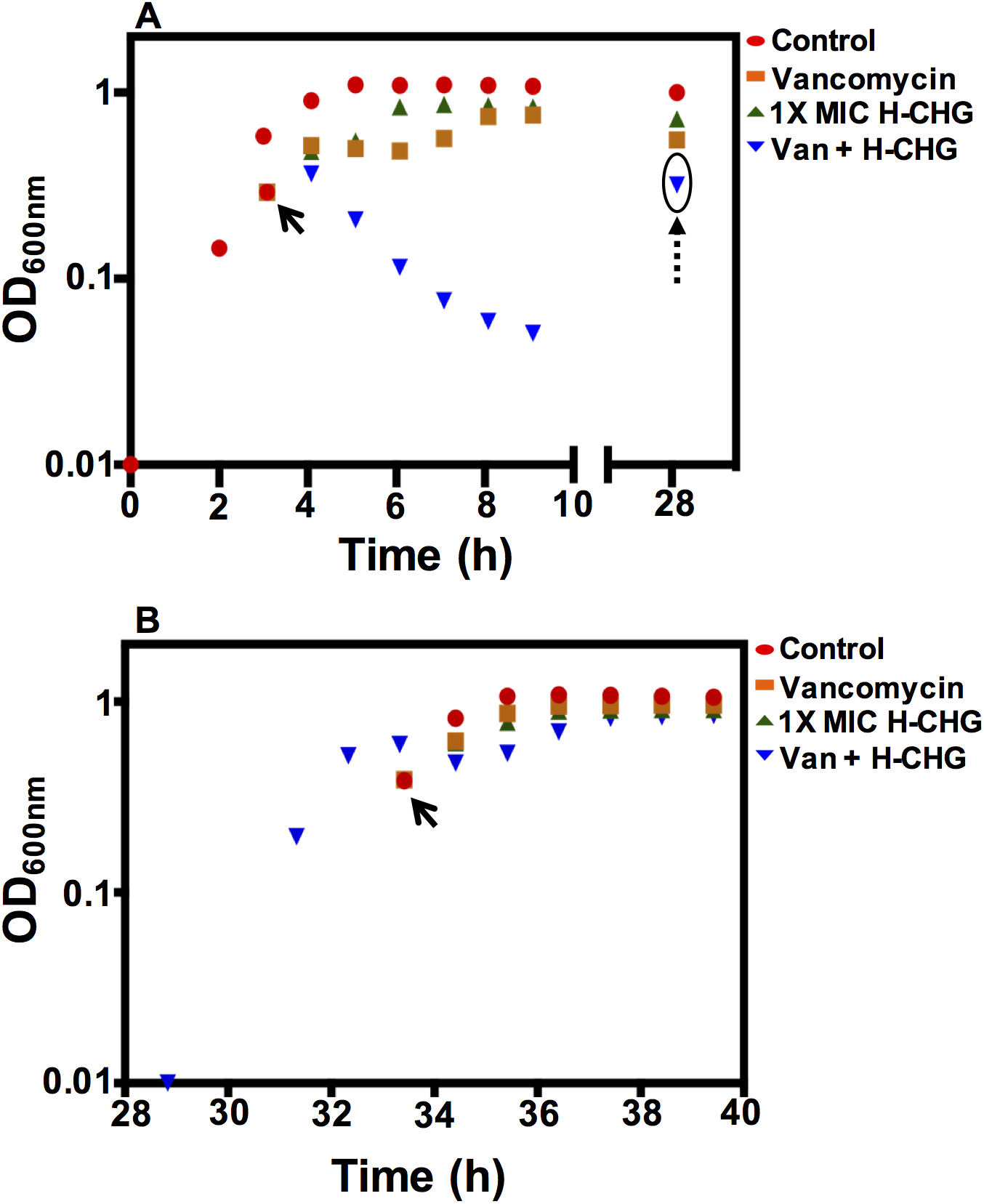
*E. faecium* 1,231,410 can adapt to vancomycin/H-CHG synergy. The growth kinetics of *E. faecium* 410 in the presence of vancomycin and H-CHG were observed over a two-day (40 h) growth curve. Panel (A) Representative OD_600_ of *E. faecium* 410 after treatment with 0X (control; red circles), vancomycin (orange squares), H-CHG (green triangles) or vancomycin and H-CHG (inverted blue triangles). *E. faecium* culture was grown at 37°C in BHI until OD_600_ reached 0.6 and equal volumes of cultures were split into BHI with different antimicrobials (shown by arrow) as described in materials and methods. OD_600_ values were monitored for 6 h and after 24 h, the vancomycin and H-CHG-treated recovered culture (circled and indicated with dashed arrow) was used as an inoculum to repeat the growth curve (shown in panel B).

The next day, the recovered culture (from the vancomycin + H-CHG growth curve) was used as inoculum to repeat the growth curve experiment (Fig. 3B). Interestingly, the growth inhibition phenotypes observed for the first growth curve experiment were not observed in this second passage. Most strikingly, cell lysis was no longer observed for the vancomycin- and chlorhexidine-treated culture. This is an important observation since it indicates that synergy mutant(s) that do not lyse in the presence of vancomycin and H-CHG can readily emerge.

### A synergy escape mutant has a mutation in *pstB*

We colony-purified a synergy escaper mutant (SE101) from the second growth curve cycle, as described in the materials and methods. The growth kinetics of SE101 in the presence of vancomycin and H-CHG confirmed that the synergism phenotype is altered in this strain (Fig. 4A and B). SE101 was initially growth-inhibited in the presence of vancomycin and H-CHG, but after 3 h, OD_600_ values began to increase, unlike what is observed for the wild-type. Significant differences in OD_600_ values were observed for SE101 compared to the wild-type for time points 3 h after addition of vancomycin and H-CHG (*P*-value < 0.05 using one-tailed Student’s *t* test).

**Figure 4.**
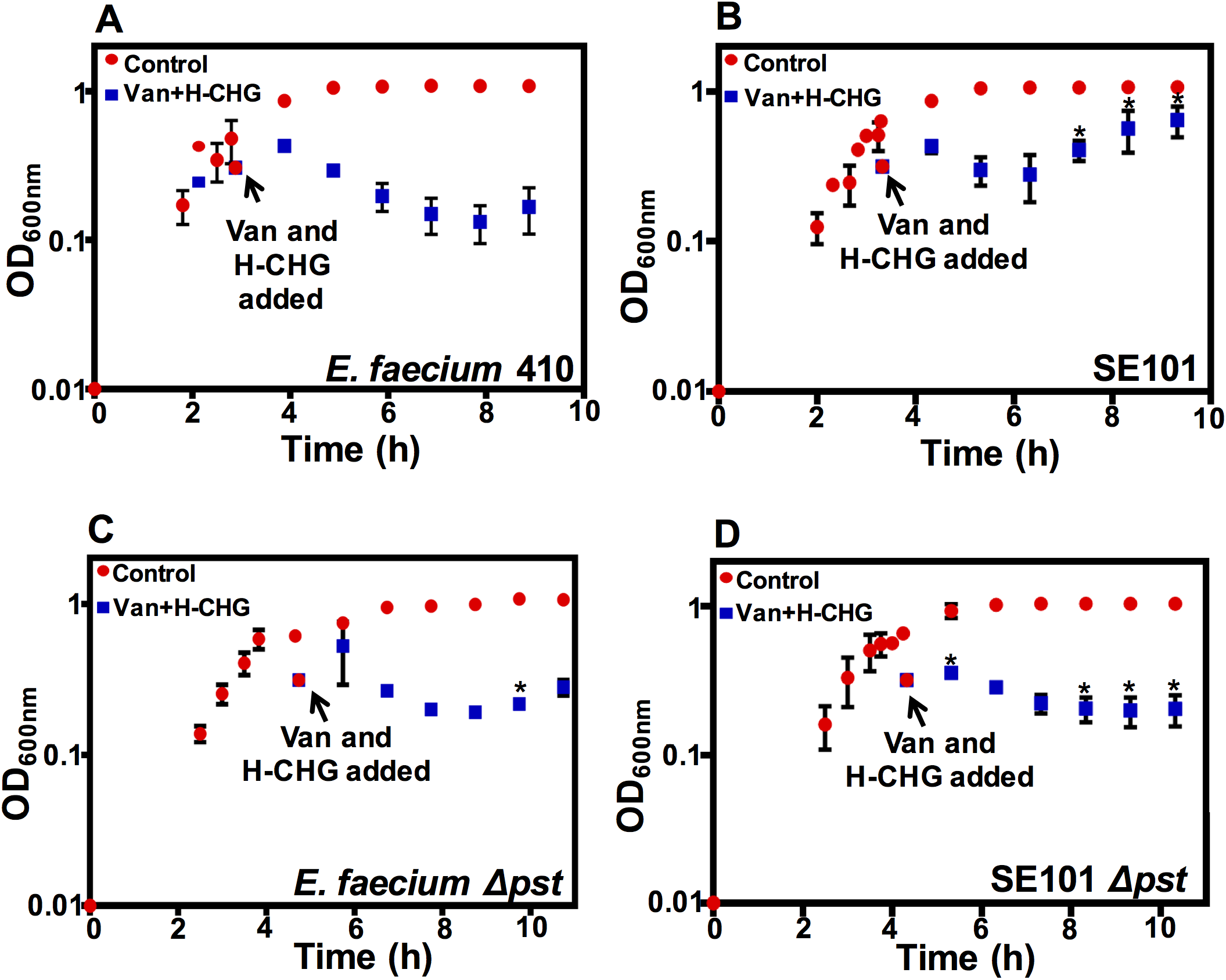
Mutations in the phosphate-specific transport (*pst*) operon result in escape from vancomycin-H-CHG synergy. Growth of (A) *E. faecium* 410 wild-type, (B) SE101, (C) *E. faecium* 410Δ*pst*, and (D) SE101Δ*pst*. *E. faecium* was cultured in BHI until the OD_600_ reached 0.6. Equal volumes of cultures were split into BHI (control; red circles) or BHI containing vancomycin (50 μg/ml) and H-CHG (4.9 μg/ml) (blue squares). OD_600_ values were monitored for 6 h and the 24 h time point was recorded. Error bars indicate standard deviations from n=3 independent experiments. Significance was assessed using the one-tailed Student’s *t*-test. * denotes *P*-value < 0.05. Stars indicate significant differences between vancomycin- and H-CHG-treated cultures in panel B versus A, in panel C versus A, and in panel D versus B.

Utilizing whole genome sequencing, we identified a mutation conferring a S199L substitution in PstB (EFTG_01173) in SE101. As a result of this substitution, the protein is predicted to fold into a beta-strand instead of a coil (48). The *pst* (phosphate-specific transport) operon has been well characterized in *E. coli* and consists of *pstSCAB* and *phoU* (a regulator). In phosphate-starvation conditions, inorganic phosphate (Pi) binds PstS and is released into the cytoplasm by the inner membrane channel formed by PstA-PstC. PstB energizes this channel by hydrolyzing ATP (49). The *pst* system and antimicrobial susceptibility has been previously linked in *E. faecium*. We identified a non-synonymous substitution in *phoU*, a negative regulator of the *pst* operon, in a chlorhexidine-adapted *E. faecium* 1,231,410 derivative that had reduced chlorhexidine and daptomycin susceptibilities and decreased intracellular Pi levels (23).

We quantified the levels of intracellular inorganic phosphate (Pi) in the wild-type strain and in SE101 under routine culture conditions. However, the levels were not significantly different for any time point assayed (Fig. S3).

To investigate the function of the *pstB* mutation further, we deleted the complete *pst* transport system (EFTG_01170-74) in SE101 and in the *E. faecium* 1,231,410 wild-type. The growth observed in the presence of vancomycin and H-CHG for the SE101Δ*pst* deletion mutant was significantly different as compared to SE101 (Fig. 4B and D; *P*-value < 0.05 using one-tailed Student’s *t* test). Specifically, unlike SE101, the OD_600_ values did not increase for the SE101Δ*pst* deletion mutant after 3 h in the presence of vancomycin and H-CHG. Conversely, deletion of the *pst* transport system from the wild-type strain did not substantially alter OD_600_ values in response vancomycin and H-CHG (Fig. 4A and C). We conclude that the *pstB* mutation in SE101 confers protection from killing by vancomycin and H-CHG co-treatment by an as yet undetermined mechanism, allowing cells to grow in the presence of the two drugs (Fig. 4B).

### Conclusions and perspective

We previously reported that *E. faecium* 1,231,410 exhibits increased susceptibility to vancomycin in the presence of chlorhexidine (19). The goal of the current study was to identify molecular contributors to this phenotype. The long-term goal is to use this information to identify less toxic compounds that could be compounded with vancomycin to exploit this vulnerability. That said, products incorporating chlorhexidine with antibiotics have been previously reported. Synergism between vancomycin and chlorhexidine was previously reported in methicillin-resistant *Staphylococcus aureus*, where chitosan-based sponges were utilized for localized delivery of these two synergistic compounds that inhibited *S. aureus* growth for 21 days (50). Another study exploited synergism between chlorhexidine and β-lactam antibiotics and synthesized hybrid organic salts (GUMBOS), which were effective against clinical isolates of Gram-positive and Gram-negative bacteria (51).

In our previous report, we proposed three models that are not mutually exclusive that could explain this phenotype. The models are reiterated here. Model 1 is that altered Pbp levels in the presence of chlorhexidine prevent D-Ala-D-Lac precursors from being cross-linked. Model 2 proposes that the chlorhexidine stress response alters substrate pools for peptidoglycan synthesis, resulting in vancomycin-sensitive termini that are neither D-Ala-D-Ala nor D-Ala-D-Lac. Finally, model 3 is that post-translational regulation of VanX and/or VanY prevents depletion of D-Ala-D-Ala termini from peptidoglycan precursors in the presence of chlorhexidine, thereby causing cells to be sensitive to vancomycin. In terms of Model 2, the results of the D-lactate amendment study (Table 3) indicate that D-Ala-D-Lac termini (and therefore, induction of the vancomycin resistance genes) are required for the synergy phenotype. Model 3 is not supported by our observation that *vanY_A_* deletion has no impact on the synergy phenotype (Fig. S1), but *vanX_A_* remains to be investigated, and therefore this model has not been fully assessed.

Growth analyses of *E. faecium* deletion mutants provide some support for Model 1. Upon *ddcP* deletion, which is predicted to result in increased availability of pentapeptide precursors for cross-linking, *E. faecium* 1,231,410 cells maintained susceptibility to the synergistic action of vancomycin and chlorhexidine, but the cells did not lyse. These results suggest that *ddcP* contributes to weakening of the cell wall in the presence of the two drugs. Our previously published transcriptomic study identified up to 5-fold induction of *ddcP* in *E. faecium* 1,231,410 cultures treated with the MIC of H-CHG for 15 minutes, as compared to untreated cultures (19). It is possible that in the presence of H-CHG, DdcP actively trims the terminal D-Ala from peptidoglycan precursors and generates tetrapeptides. At the same time, in the presence of vancomycin, the combined activities of the vancomycin resistance genes result in pentapeptides terminating in D-Lac. The relative availability of penta- and tetrapeptides with chemically different termini likely impacts the overall efficiency of cross-linking and the strength of the cell wall. Since all Pbps do not have the same affinity for tetra-versus pentapeptides, unacceptable precursors for transpeptidation are synthesized in the presence of both vancomycin and chlorhexidine, and the cells lyse. However, complicating this explanation, we did not observe any difference in growth phenotype between the L,D transpeptidase (*ldt_fm_*) deletion mutant and the wild-type in the presence of vancomycin and H-CHG (Fig. S1F and G). As reported previously, activity of Ldt_fm_ is dependent on availability of tetrapeptides (42, 43). If (and how) Ldt_fm_ activity changes in the presence of vancomycin and H-CHG remains to be elucidated. A critical set of future experiments that are relevant to both Models 1 and 2 will be to analyze cytoplasmic peptidoglycan precursor pools and mature peptidoglycan structures in *E. faecium* wild-type and *ddcP* mutant cultures exposed to vancomycin and chlorhexidine. This analysis would allow us to analyze the relative balance of tetra-versus pentapeptide termini, as well as their chemical compositions. Moreover, more detailed assessments of cell viability and structure would also be useful, including live/dead staining and electron microscopy.

We also found that synergy escaper mutants (i.e., cells that failed to lyse) arose after 24 h of exposure to both vancomycin and chlorhexidine. A spontaneous non-synonymous substitution in *pstB* conferred a survival advantage in the presence of the two antimicrobials. However, the exact mechanism(s) of how the Pst system impacts antimicrobial susceptibility in *E. faecium* is unknown. Critical experiments for the future are to analyze Pi levels under stressed conditions (i.e., in the presence of vancomycin and chlorhexidine), and to perform analysis of cytoplasmic peptidoglycan precursor pools and mature peptidoglycan structures, as described above.

Overall, our study highlights the complexity of the enterococcal cell wall stress response in response to combination antimicrobial therapy and identifies a novel contributor (*pstB*) to this response. We additionally present a collection of deletion mutants, validated by genome sequencing, that are of use for future studies of *E. faecium* cell wall biology.

## Acknowledgements

This work was supported by start-up funds from the University of Texas at Dallas and the Cecil H. and Ida Green Chair in Systems Biology Science to K.L.P. We are grateful to Dr. Breck Duerkop for providing the plasmid pLT06-*tet*.

## Supplemental Figures and Tables

**Figure S1.**
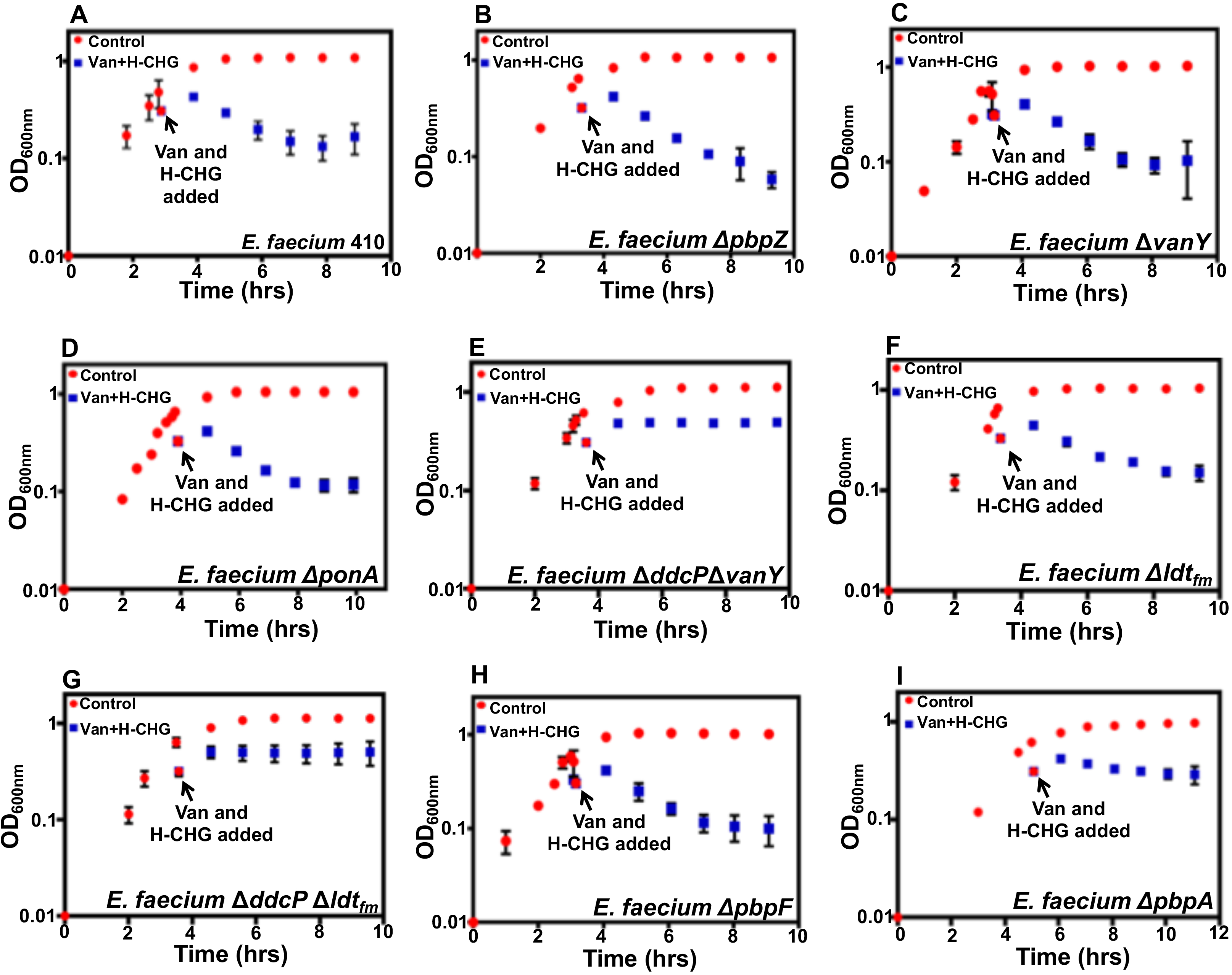
Representative optical density (OD_600_) of (A) *E. faecium* 410 wild-type, (B) Δ*pbpZ*, (C) Δ*vanY*, (D) Δ*ponA*, (E) Δ*ddcP* Δ*vanY*, (F) Δ*ldt_fm_*, (G) Δ*ddcP* Δ*ldt_fm_*, (H) Δ*pbpF*, and (I) Δ*pbpA* after vancomycin and chlorhexidine treatment. *E. faecium* was cultured at 37°C in BHI broth until the OD_600_ reached 0.6 as described in the materials and methods. Equal volumes of culture were split into BHI containing 0X (control; red circles) or vancomycin and chlorhexidine (Van and H-CHG; blue squares). OD_600_ values were monitored for 6 h. Error bars indicate standard deviations from three independent experiments.

**Figure S2.**
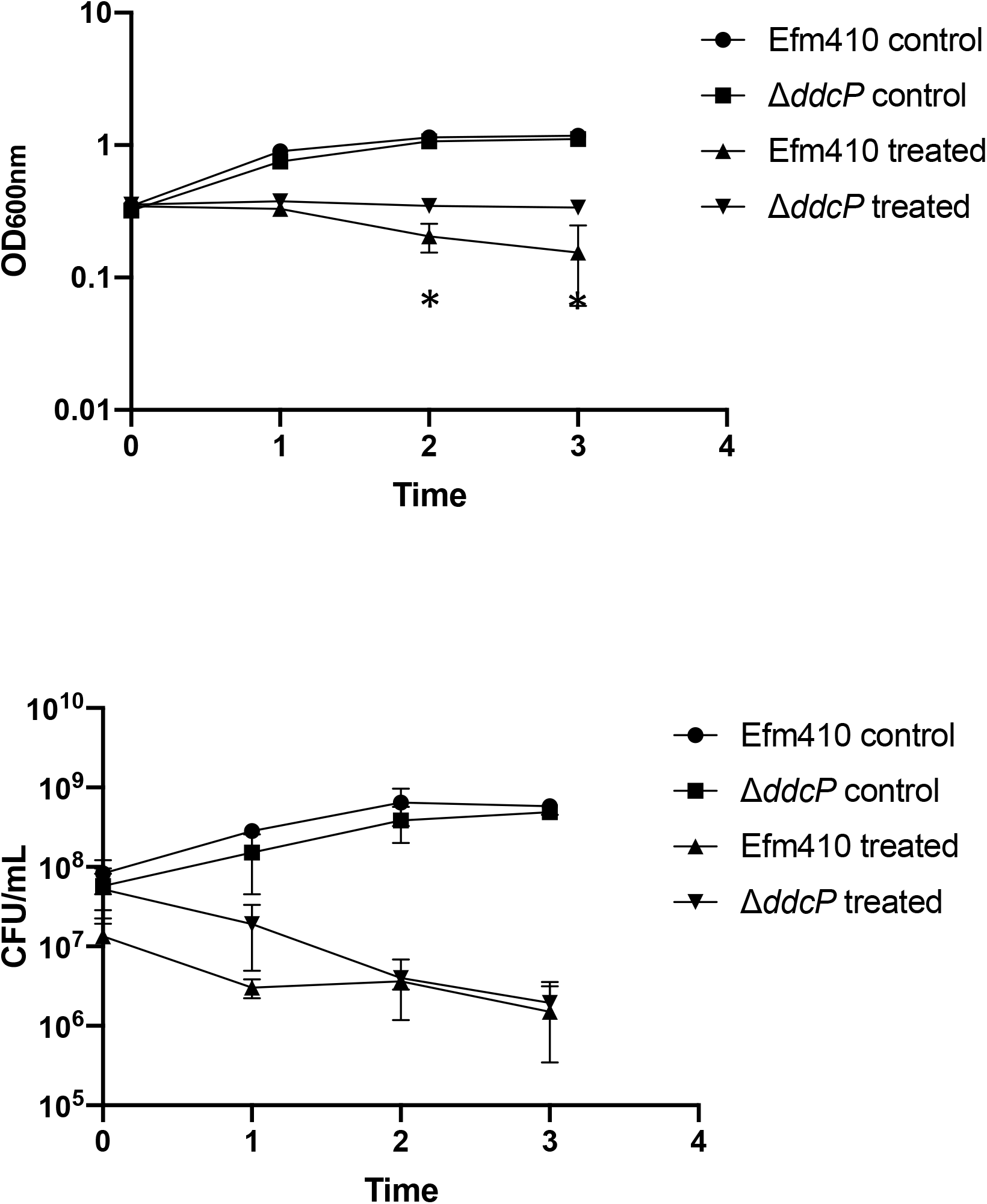
A Δ*ddcP* mutant dies but does not lyse in the presence of vancomycin and H-CHG. Optical density (OD_600nm_) (A) and CFU/mL (B) of *E. faecium* 1,231,410 wild-type (*E. faecium* 410) and the *ddcP* deletion mutant with (“treated”) and without (“control”) vancomycin and H-CHG treatment. *E. faecium* was cultured in BHI broth until the OD_600_ reached 0.6, as described in materials and methods. Equal volumes of cultures were split into BHI or BHI containing vancomycin (50 μg/ml) and H-CHG (4.9 μg/ml). OD_600_ values and CFU/mL were monitored for 3 h. Error bars indicate standard deviations from n=3 independent experiments. Significance was assessed using the one-tailed Student’s *t*-test. * denotes *P*-value < 0.05. Stars indicate significant differences between vancomycin- and H-CHG-treated cultures.

**Figure S3.**
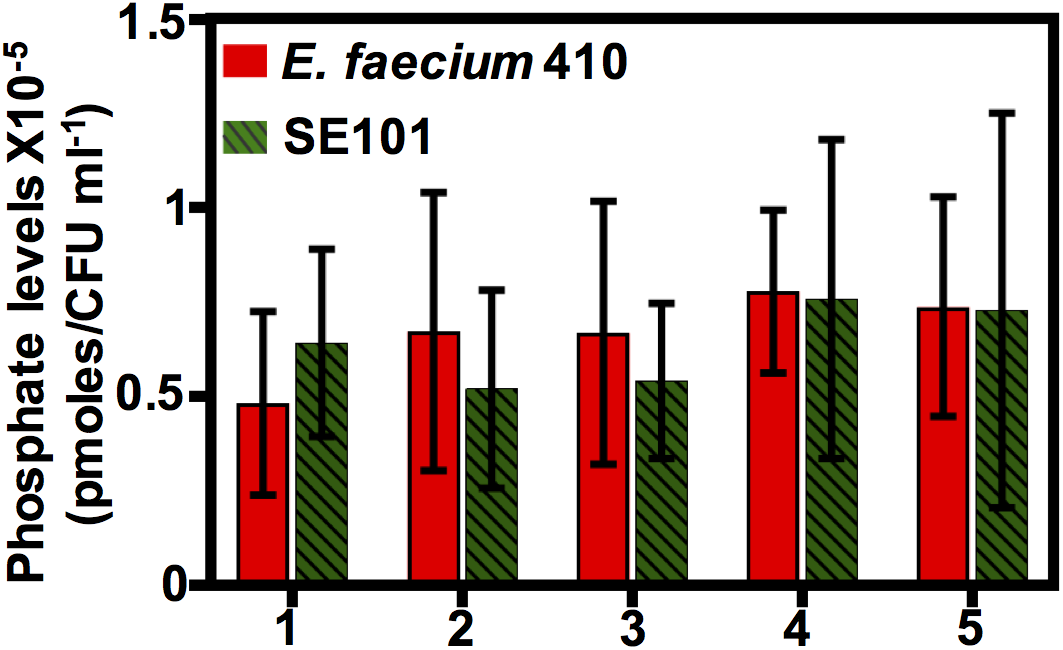
Quantification of intracellular organic phosphate (Pi) levels in *E. faecium* 1,231,410 wild-type and SE101 synergy escaper mutant. Intracellular Pi levels were measured for both strains at different growth time points (OD_600_ 0.4-1.0) as described in materials and methods. The levels (pmoles) were normalized using CFU count. Standard deviation was calculated from n=5 independent experiments and significance value was calculated using one-tailed Student’s *t* test. Time points: 1, OD_600_ 0.4-0.5; 2, OD_600_ 0.6-0.7; 3, OD_600_ 0.7-0.8; 4, OD_600_ 0.8-0.9; OD_600_ 1.0-1.5.

**Table S1.**
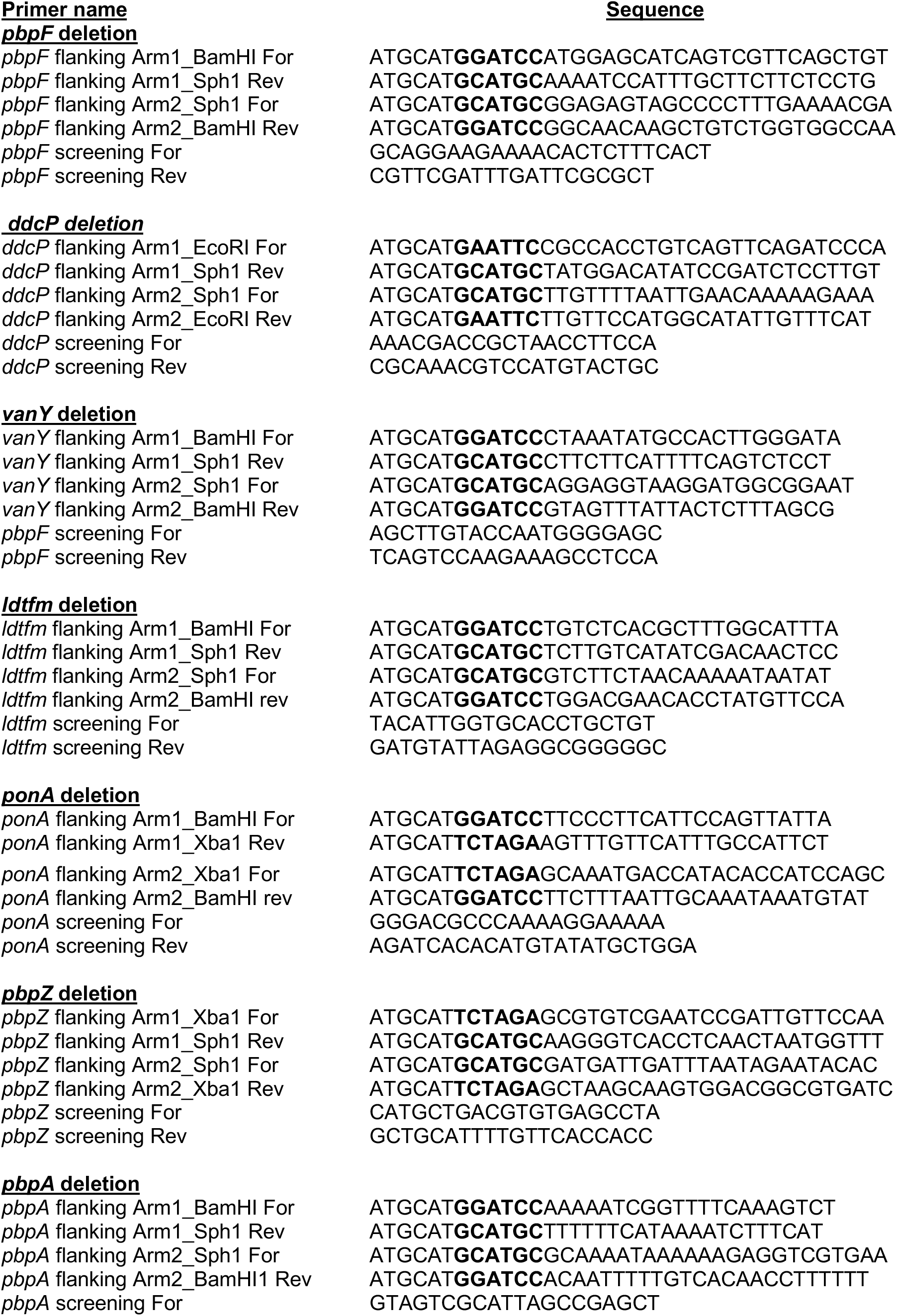
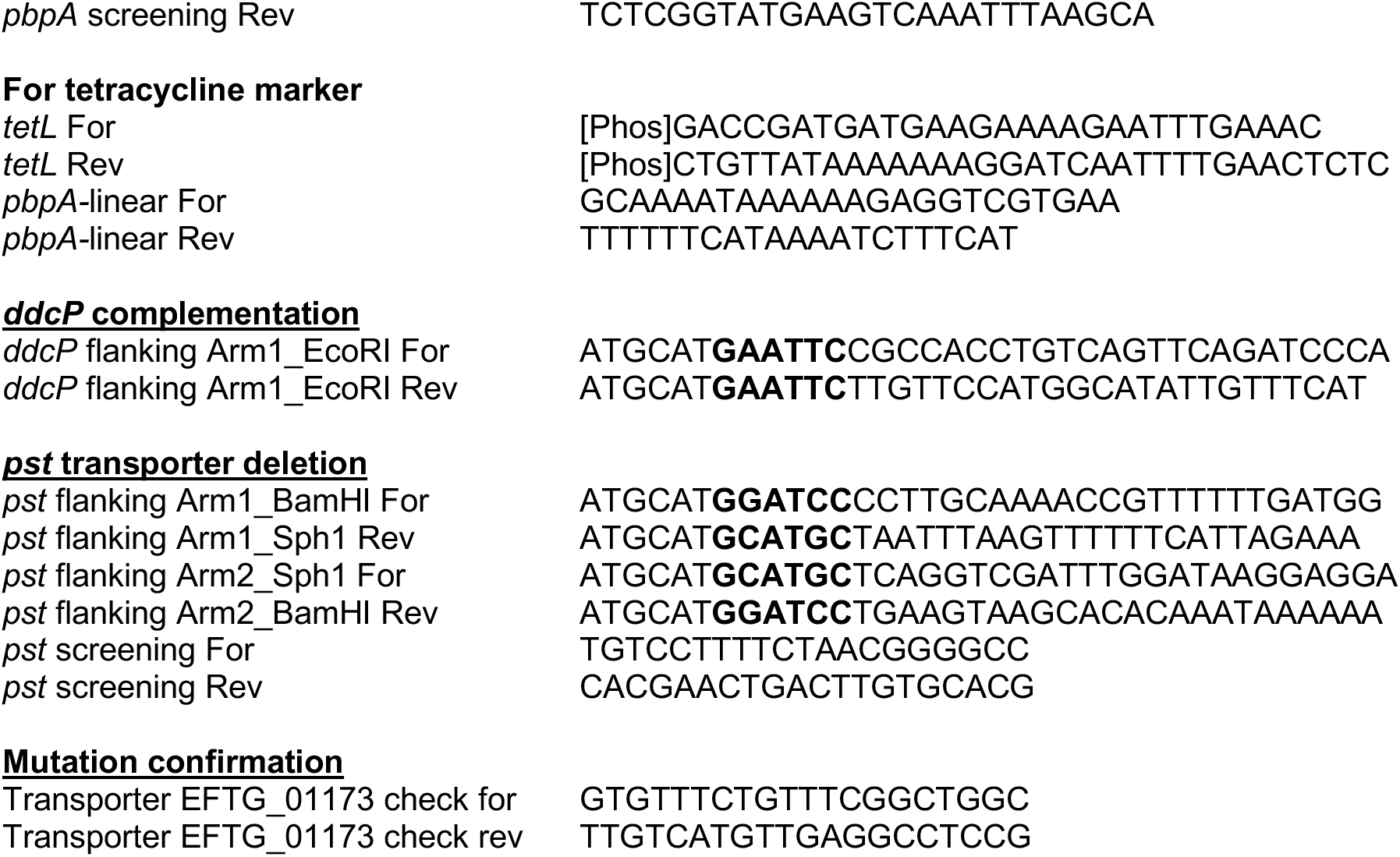
List of primers used in the study.

## Notes

### Competing Interest Statement

The authors have declared no competing interest.

### Summary of Updates

(1) We performed genome sequencing to confirm gene deletions described in the study. Of the mutants described in the previous version, three could not be confirmed. They have been removed from the manuscript. The rest of the mutants are confirmed. The sequence data have been deposited in the Sequence Read Archive. (2) Additional experiments regarding the impact of ddcP deletion on E. faecium 1,231,410 have been performed.

## References

1. Agudelo Higuita NI, Huycke MM. 2014. Enterococcal Disease, Epidemiology, and Implications for Treatment. In: Gilmore MS, Clewell DB, Ike Y, Shankar N, editors. Enterococci: From Commensals to Leading Causes of Drug Resistant Infection. Boston.

2. Lebreton F, Manson AL, Saavedra JT, Straub TJ, Earl AM, Gilmore MS. Tracing the Enterococci from Paleozoic Origins to the Hospital. Cell. 2017;169(5):849–61 e13.

3. Reynolds PE. Structure, biochemistry and mechanism of action of glycopeptide antibiotics. European Journal of Clinical Microbiology & Infectious Diseases. 1989;8(11):943–50.

4. Courvalin P. Vancomycin resistance in gram-positive cocci. Clinical Infectious Diseases. 2006;42 Suppl 1:S25–34.

5. Handwerger S, Pucci MJ, Volk KJ, Liu J, Lee MS. The cytoplasmic peptidoglycan precursor of vancomycin-resistant *Enterococcus faecalis* terminates in lactate. Journal of Bacteriology. 1992;174(18):5982–4.

6. Allen NE, Hobbs JN, Jr., Richardson JM, Riggin RM. Biosynthesis of modified peptidoglycan precursors by vancomycin-resistant *Enterococcus faecium*. FEMS Microbiology Letters. 1992;77(1-3):109–15.

7. Messer J, Reynolds PE. Modified peptidoglycan precursors produced by glycopeptide-resistant enterococci. FEMS Microbiology Letters. 1992;73(1-2):195–200.

8. Liu J, Volk KJ, Lee MS, Pucci M, Handwerger S. Binding studies of vancomycin to the cytoplasmic peptidoglycan precursors by affinity capillary electrophoresis. Analytical Chemistry. 1994;66(14):2412–6.

9. Leclercq R, Derlot E, Duval J, Courvalin P. Plasmid-mediated resistance to vancomycin and teicoplanin in *Enterococcus faecium*. The New England Journal of Medicine. 1988;319(3):157–61.

10. Woodford N. Glycopeptide-resistant enterococci: a decade of experience. Journal of Medical Microbiology. 1998;47(10):849–62.

11. Munoz-Price LS, Hota B, Stemer A, Weinstein RA. Prevention of bloodstream infections by use of daily chlorhexidine baths for patients at a long-term acute care hospital. Infection Control and Hospital Epidemiology. 2009;30(11):1031–5.

12. Muto CA, Jernigan JA, Ostrowsky BE, Richet HM, Jarvis WR, Boyce JM, et al. SHEA guideline for preventing nosocomial transmission of multidrug-resistant strains of *Staphylococcus aureus* and *Enterococcus*. Infection Control and Hospital Epidemiology. 2003;24(5):362–86.

13. LeDell K, Muto CA, Jarvis WR, Farr BM. SHEA guideline for preventing nosocomial transmission of multidrug-resistant strains of *Staphylococcus aureus* and *Enterococcus*. Infection Control and Hospital Epidemiology. 2003;24(9):639–41.

14. Komljenovic I, Marquardt D, Harroun TA, Sternin E. Location of chlorhexidine in DMPC model membranes: a neutron diffraction study. Chemistry and Physics of Lipids. 2010;163(6):480–7.

15. Koontongkaew S, Jitpukdeebodintra S. Interaction of chlorhexidine with cytoplasmic membranes of *Streptococcus mutans* GS-5. Caries Research. 1995;29(5):413–7.

16. Hugo WB, Longworth AR. Some Aspects of the Mode of Action of Chlorhexidine. The Journal of Pharmacy and Pharmacology. 1964;16:655–62.

17. Hugo WB, Longworth AR. Cytological Aspects of the Mode of Action of Chlorhexidine Diacetate. The Journal of Pharmacy and Pharmacology. 1965;17:28–32.

18. Hugo WB, Longworth AR. The effect of chlorhexidine on the electrophoretic mobility, cytoplasmic constituents, dehydrogenase activity and cell walls of *Escherichia coli* and *Staphylococcus aureus*. The Journal of Pharmacy and Pharmacology. 1966;18(9):569–78.

19. Bhardwaj P, Ziegler E, Palmer KL. Chlorhexidine Induces VanA-Type Vancomycin Resistance Genes in Enterococci. Antimicrobial Agents and Chemotherapy. 2016;60(4):2209–21

20. Palmer KL, Godfrey P, Griggs A, Kos VN, Zucker J, Desjardins C, et al. Comparative genomics of enterococci: variation in *Enterococcus faecalis*, clade structure in *E. faecium*, and defining characteristics of *E. gallinarum* and *E. casseliflavus*. MBio. 2012;3(1):e00318–11.

21. Adams HM, Li X, Mascio C, Chesnel L, Palmer KL. Mutations associated with reduced surotomycin susceptibility in *Clostridium difficile* and *Enterococcus* species. Antimicrobial Agents and Chemotherapy. 2015;59(7):4139–47.

22. Thurlow LR, Thomas VC, Hancock LE. Capsular polysaccharide production in *Enterococcus faecalis* and contribution of CpsF to capsule serospecificity. Journal of Bacteriology. 2009;191(20):6203–10.

23. Bhardwaj P, Hans A, Ruikar K, Guan Z, Palmer KL. Reduced Chlorhexidine and Daptomycin Susceptibility in Vancomycin-Resistant *Enterococcus faecium* after Serial Chlorhexidine Exposure. Antimicrobial Agents and Chemotherapy. 2018;62(1).

24. Mechler L, Herbig A, Paprotka K, Fraunholz M, Nieselt K, Bertram R. A novel point mutation promotes growth phase-dependent daptomycin tolerance in *Staphylococcus aureus*. Antimicrobial Agents and Chemotherapy. 2015;59(9):5366–76.

25. Zarlenga LJ, Gilmore MS, Sahm DF. Effects of amino acids on expression of enterococcal vancomycin resistance. Antimicrobial Agents and Chemotherapy. 1992;36(4):902–5.

26. van der Aart LT, Lemmens N, van Wamel WJ, van Wezel GP. Substrate Inhibition of VanA by d-Alanine Reduces Vancomycin Resistance in a VanX-Dependent Manner. Antimicrobial Agents and Chemotherapy. 2016;60(8):4930–9.

27. Hancock LE, Murray BE, Sillanpaa J. 2014. Enterococcal Cell Wall Components and Structures. In: Gilmore MS, Clewell DB, Ike Y, Shankar N, editors. Enterococci: From Commensals to Leading Causes of Drug Resistant Infection. Boston.

28. Coyette J, Hancock L. Enterococcal Cell Wall. In: Gilmore MS, editor. The Enterococci: Pathogenesis, Molecular Biology, and Antibiotic Resistance. Washington, D.C.: ASM Press; 2002. p. 385–408.

29. Ghuysen JM. Use of bacteriolytic enzymes in determination of wall structure and their role in cell metabolism. Bacteriol Rev. 1968;32(4 Pt 2):425–64.

30. Schleifer KH, Kandler O. Peptidoglycan types of bacterial cell walls and their taxonomic implications. Bacteriol Rev. 1972;36(4):407–77.

31. Navarre WW, Schneewind O. Surface proteins of gram-positive bacteria and mechanisms of their targeting to the cell wall envelope. Microbiology and Molecular Biology Reviews. 1999;63(1):174–229.

32. Vollmer W, Blanot D, de Pedro MA. Peptidoglycan structure and architecture. FEMS Microbiology Reviews. 2008;32(2):149–67.

33. van Heijenoort J. Assembly of the monomer unit of bacterial peptidoglycan. Cellular and Molecular Life Sciences. 1998;54(4):300–4.

34. Anderson JS, Matsuhashi M, Haskin MA, Strominger JL. Lipid-Phosphoacetylmuramyl-Pentapeptide and Lipid-Phosphodisaccharide-Pentapeptide: Presumed Membrane Transport Intermediates in Cell Wall Synthesis. Proceedings of the National Academy of Sciences of the United States of America. 1965;53:881–9.

35. Anderson JS, Strominger JL. Isolation and utilization of phospholipid intermediates in cell wall glycopeptide synthesis. Biochemical and Biophysical Research Communications. 1965;21(5):516–21.

36. van Dam V, Sijbrandi R, Kol M, Swiezewska E, de Kruijff B, Breukink E. Transmembrane transport of peptidoglycan precursors across model and bacterial membranes. Mol Microbiol. 2007;64(4):1105–14.

37. van Heijenoort J. Formation of the glycan chains in the synthesis of bacterial peptidoglycan. Glycobiology. 2001;11(3):25R–36R.

38. Scheffers DJ, Pinho MG. Bacterial cell wall synthesis: new insights from localization studies. Microbiology and Molecular Biology Reviews. 2005;69(4):585–607.

39. Rice LB, Carias LL, Rudin S, Hutton R, Marshall S, Hassan M, et al. Role of class A penicillin-binding proteins in the expression of beta-lactam resistance in *Enterococcus faecium*. Journal of Bacteriology. 2009;191(11):3649–56.

40. Ghuysen JM. Serine beta-lactamases and penicillin-binding proteins. Annual Review of Microbiology. 1991;45:37–67.

41. Goffin C, Ghuysen JM. Multimodular penicillin-binding proteins: an enigmatic family of orthologs and paralogs. Microbiology and Molecular Biology Reviews. 1998;62(4):1079–93.

42. Mainardi JL, Fourgeaud M, Hugonnet JE, Dubost L, Brouard JP, Ouazzani J, et al. A novel peptidoglycan cross-linking enzyme for a beta-lactam-resistant transpeptidation pathway. The Journal of Biological Chemistry. 2005;280(46):38146–52.

43. Mainardi JL, Morel V, Fourgeaud M, Cremniter J, Blanot D, Legrand R, et al. Balance between two transpeptidation mechanisms determines the expression of beta-lactam resistance in *Enterococcus faecium*. The Journal of Biological Chemistry. 2002;277(39):35801–7.

44. Mainardi JL, Legrand R, Arthur M, Schoot B, van Heijenoort J, Gutmann L. Novel mechanism of beta-lactam resistance due to bypass of DD-transpeptidation in *Enterococcus faecium*. The Journal of Biological Chemistry. 2000;275(22):16490–6.

45. Kristich CJ, Djoric D, Little JL. Genetic basis for vancomycin-enhanced cephalosporin susceptibility in vancomycin-resistant enterococci revealed using counterselection with dominant-negative thymidylate synthase. Antimicrobial Agents and Chemotherapy. 2014;58(3):1556–64.

46. Nicas TI, Wu CY, Hobbs JN, Jr., Preston DA, Allen NE. Characterization of vancomycin resistance in *Enterococcus faecium* and *Enterococcus faecalis*. Antimicrobial Agents and Chemotherapy. 1989;33(7):1121–4.

47. Williamson R, Al-Obeid S, Shlaes JH, Goldstein FW, Shlaes DM. Inducible resistance to vancomycin in *Enterococcus faecium* D366. The Journal of Infectious Diseases. 1989;159(6):1095–104.

48. Rice P, Longden I, Bleasby A. EMBOSS: the European Molecular Biology Open Software Suite. Trends Genet. 2000;16(6):276–7.

49. Chan FY, Torriani A. PstB protein of the phosphate-specific transport system of *Escherichia coli* is an ATPase. Journal of Bacteriology. 1996;178(13):3974–7.

50. Smith JK, Moshref AR, Jennings JA, Courtney HS, Haggard WO. Chitosan sponges for local synergistic infection therapy: a pilot study. Clin Orthop Relat Res. 2013;471(10):3158–64.

51. Cole MR, Hobden JA, Warner IM. Recycling antibiotics into GUMBOS: a new combination strategy to combat multi-drug-resistant bacteria. Molecules. 2015;20(4):6466–87.

